# Protein function achieved through multiple covalent states

**DOI:** 10.1101/828368

**Authors:** Diego Butera, Philip J. Hogg

## Abstract

The structure of proteins is defined by two main types of covalent bonds; the peptide bonds that link the amino acid residues and disulfide bonds that link pairs of cysteine amino acids. Disulfide bonds are introduced during protein folding and their formation is assumed to be complete in the mature, functional protein. We tested this assumption by quantifying the redox state of disulfide bonds in human blood proteins in their native environment. Using a differential cysteine alkylation and mass spectrometry method, we measured the redox state of disulfide bonds in circulating fibrinogen and von Willebrand factor. There is an extraordinary disulfide lability in the proteins, with 27 bonds in the two proteins ranging from 3 to 50% reduced in healthy human donors. Modelling of the data indicates that the proteins exist in hundreds of different disulfide-bonded states in the circulation. Different covalent states of fibrinogen are associated with different binding activities and their distribution is changed by fluid shear forces and altered in patients with cardiovascular disease, indicating that the different states have different functions and are dynamic. These findings have implications for protein function generally and how proteins are targeted in experimental settings and for therapeutic purposes.

## Introduction

Protein Data Bank structures currently contain more than 180,000 disulfide bonds^1^. In human proteins, these bonds are found mostly in membrane proteins (plasma and organelle) and proteins that are secreted by cells. Although, a growing number of disulfide bonds are being identified in proteins that function in the nucleus or cytoplasm. Acquisition of disulfide bonds contributed significantly to the evolution of vertebrates^2^. Most disulfides in human proteins were acquired in vertebrate ancestors and very often coincided with formation of a new protein. These bonds were almost never lost as the protein evolved and are continuing to be acquired today^3, 4^.

Disulfide bonds form during maturation of proteins in the cell and influence protein folding in ways that are not completely understood. Three different roles for these bonds in protein folding have been proposed: one where disulfides drive global protein folding^5^, another where these bonds direct protein folding in a stepwise fashion^6, 7, 8^, and a third where disulfide bond formations stabilize the native conformation rather than facilitate folding^9, 10, 11^. Thousands of disulfide-bonded protein crystal structures have revealed that these bonds are almost invariably intact.

Protein crystallization is a very exacting type of protein purification and favors the most stable, lowest energy forms of a protein. This will typically be proteins where all disulfide bonds are intact. We tested the hypothesis that proteins in their native environment may exist in states where disulfides are incompletely formed.

## Results

### Fibrinogen exists in multiple disulfide-bonded states in the circulation

Isolating a protein that contains closely spaced cysteine thiols and storing it in ambient oxygen can result in oxidation of the thiols to a disulfide bond. To exclude this possibility, we froze the redox state of cysteine thiols before the proteins were removed from their native environment, which in this case was blood plasma.

Blood from healthy donors was drawn by venipuncture into citrate as anti-coagulant, plasma prepared by centrifugation and proteins collected on antibody-coated magnetic beads. The unpaired cysteine thiols in bead-bound protein were alkylated with 2-iodo-N-phenylacetamide (^12^C-IPA), the protein isolated by SDS-PAGE, the disulfide bonds reduced with dithiothreitol, and the disulfide-bonded cysteine thiols alkylated with a stable carbon-13 isotope of IPA (^13^C-IPA)^12^. The protein was digested with trypsin and peptides quantified by HPLC and identity established by mass spectrometry. The levels of the different redox forms of the cysteines was determined from the relative abundance of peptides labelled with ^12^C-IPA and/or ^13^C-IPA (Fig. 1A and B). For more details, see Supplementary Information.

**Fig. 1.**
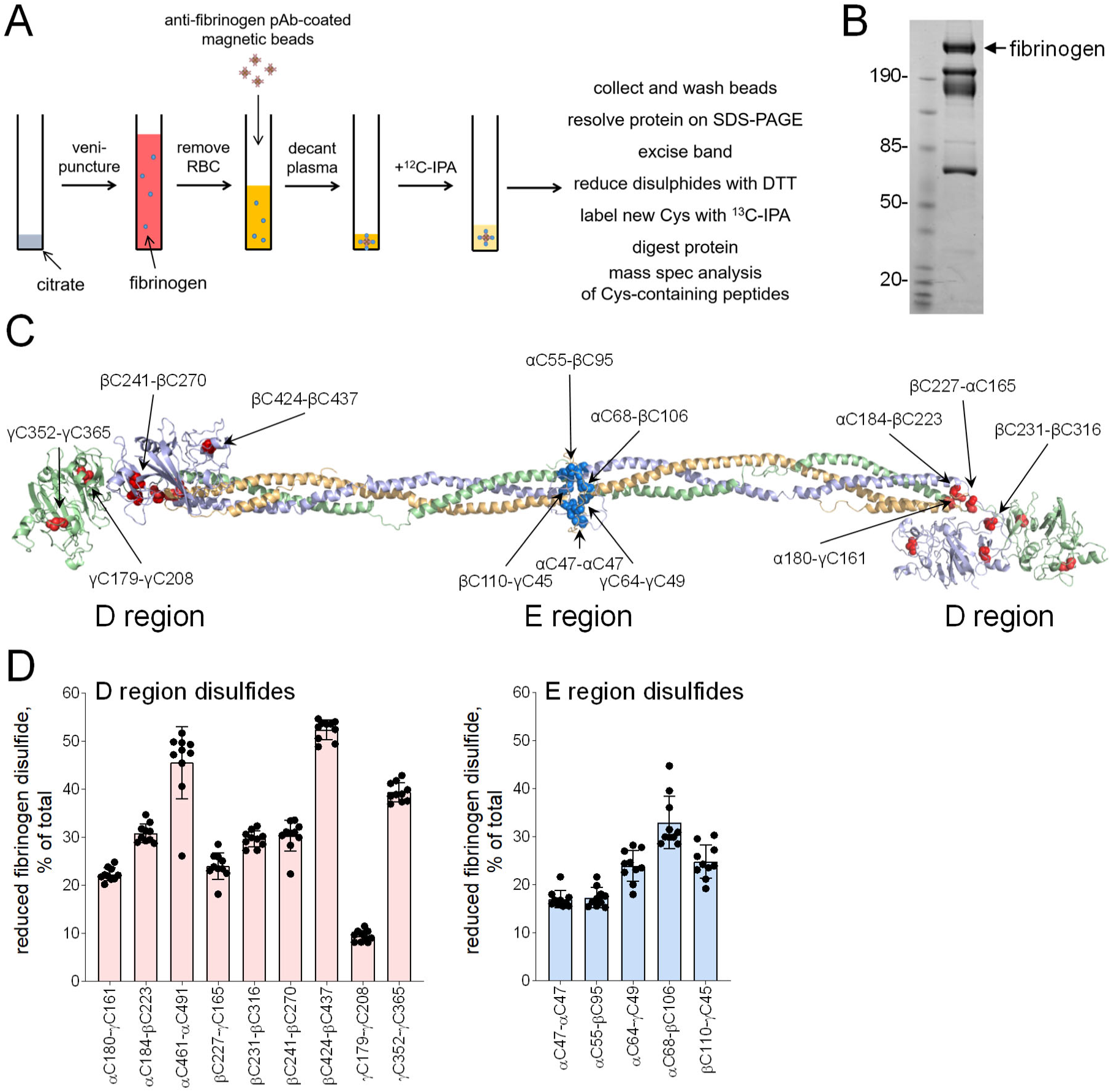
Fibrinogen exists in multiple disulfide-bonded states in the circulation. **A**. Blood from healthy donors was drawn by venipuncture into citrate as anti-coagulant, plasma prepared by centrifugation and fibrinogen collected on antibody-coated magnetic beads. The unpaired cysteine thiols in bead-bound fibrinogen was alkylated with ^12^C-IPA, the protein resolved on SDS-PAGE and the disulfide-bonded cysteine thiols alkylated with ^13^C-IPA following reduction with DTT. The protein was digested with trypsin, 17 peptides (Table S1) encompassing cysteines representing 14 of the 17 fibrinogen disulfide bonds were analyzed by HPLC and mass spectrometry and the redox state of the disulfides quantified. **B**. Example of ^12^C-IPA-labelled fibrinogen resolved on SDS-PAGE. Molecular mass standards are in the left-hand lane. **C**. Ribbon structure of fibrinogen^13^ and the positions of the 14 disulfide bonds (red spheres for the D region disulfides and cyan spheres for the E region bonds) that were mapped. The α chains are in wheat, β chains in light blue and γ chains in light green. **D**. Redox states of the nine D region and five E region disulfides in ten healthy human donors. The bars and errors are mean ± SD.

The blood protein we analysed initially was fibrinogen. Fibrinogen plays a critical role in arresting bleeding (haemostasis) and obstruction of blood vessels (thrombosis) by bridging platelets at sites of vessel injury. It was selected because of its relative abundance in blood and its well defined quaternary structure^13^ that contains 17 disulfide bonds. Fibrinogen consists of three pairs of polypeptide chains; two Aα, two Bβ and two γ chains. Two AαBβγ units are linked head-to-head to form a structure with a central E region and two peripheral D regions. The disulfide bonds in the protein are located in the globular E (7 bonds) and D regions (10 bonds). The redox state of 14 of the 17 fibrinogen disulfides was quantified; 5 in the E region and 9 bonds in the D region (Fig. 1C). Of the 14 bonds that were measured, all have been structurally defined except the D region intrachain αC461-αC491 disulfide. Remarkably, all 14 disulfide bonds exist in bound (oxidized) or cleaved (reduced) forms in the fibrinogen populations of ten healthy human donors (5 male, 5 female, 22-58 years old) (Fig. 1D). The bonds ranged from 10 to 50% reduced and there was little donor-to-donor variation.

### Number of fibrinogen covalent states is a function of conditional formation of disulfide bonds

The results indicate that circulating fibrinogen exists in multiple disulfide-bonded states. A protein containing *n* disulfide bonds, where the bonds are either formed or broken, has 2^*n*^ possible disulfide-bonded states. In the case of the 14 fibrinogen disulfides that were measured, this equates to 16,384 possible disulfide-bonded states of the protein. Inspection of the data, though, indicates that this is an overestimation of the number of species. We modelled the redox states of the five disulfide bonds in the D region β-nodule (Fig. 2A) to illustrate this point. These five bonds exist in a defined globular domain (Fig. 2B), and while their redox state is likely influenced by bond formation elsewhere in the molecule, it was assumed for modelling purposes that formation of the disulfides in the β-nodule is dependent on bonds only in that domain. Three of the five β-nodule disulfides are ~30% reduced (αC184-βC223, βC231-βC316 and βC241-βC270), while one is ~20% (βC227-γC165) and the other ~50% (βC424-βC437) reduced (Fig. 2A).

**Fig. 2.**
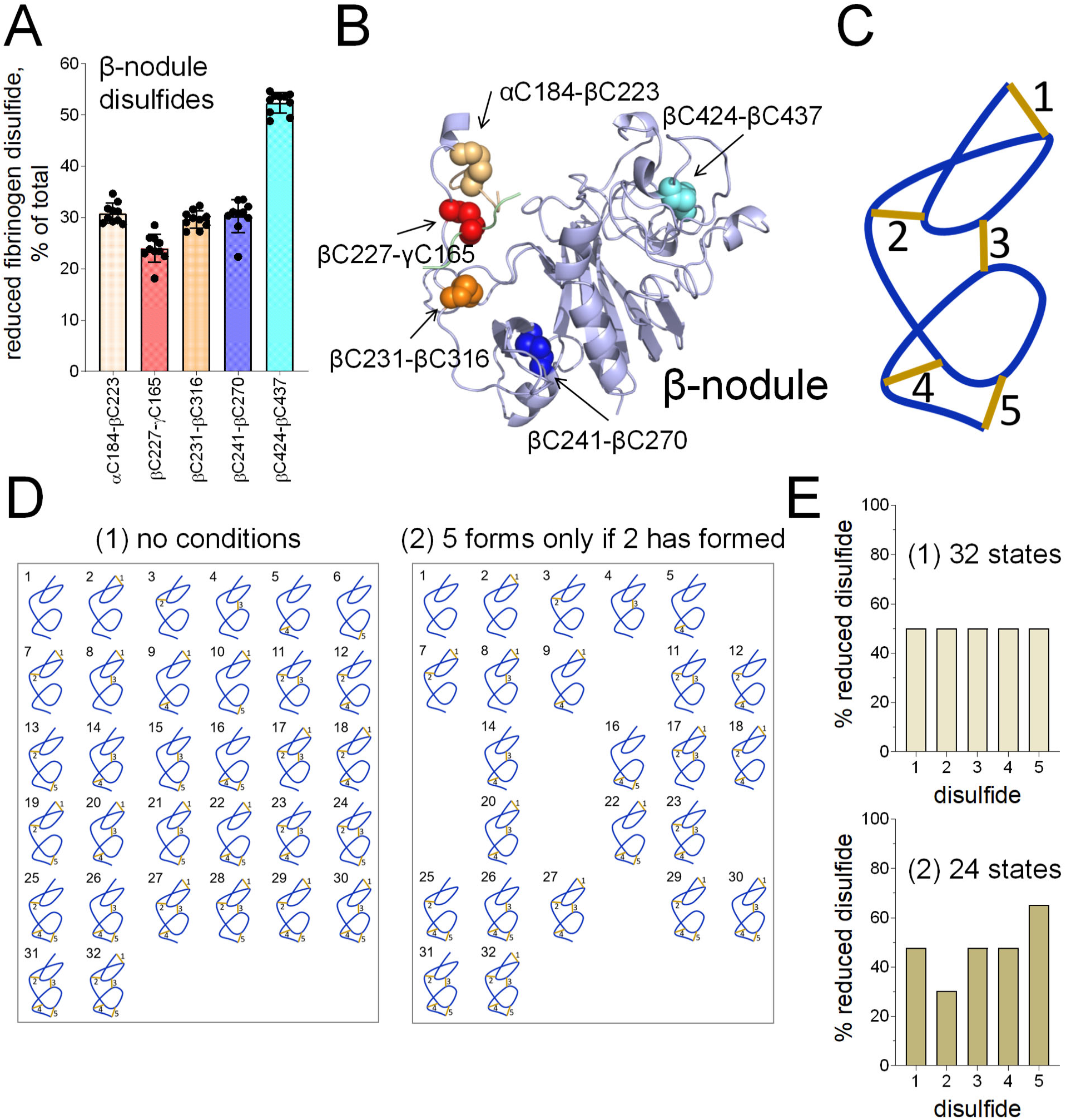
Number of fibrinogen covalent states is a function of conditional formation of disulfide bonds. **A**. Redox state of the fibrinogen β-nodules in 10 healthy human donors taken from Fig. 1D. **B**. Ribbon structure of the fibrinogen β-nodule^13^ showing the five disulfide bonds (spheres). The disulfide colour matches the coding in part A. **C**. A cartoon polypeptide containing 5 disulfide bonds used for the modelling in parts D-F. **D** and **E**. A polypeptide containing 5 disulfide bonds, where the bonds are either formed or broken, can exist in 32 possible disulfide states (part D). Assuming no conditions on disulfide formation, the 5 bonds are predicted to exist in equal proportions of reduced and oxidized states (part E). If the condition is applied that disulfide 5 only forms if disulfide 2 has formed, the total number of disulfide states reduces from 32 to 24 (part D). The pattern of the predicted redox states of the five bonds (part E) resembles the experimentally determined pattern for the five fibrinogen β-nodule disulfides (part B).

A polypeptide containing 5 disulfide bonds (Fig. 2C), where the bonds are either formed or broken, can exist in 32 (2^5^) possible disulfide-bonded states. These different states are represented in cartoon form in Fig. 2D. The reduced species for each individual disulfide was summed and this number expressed as a percentage of the total number of disulfide-bonded states to mirror the experimental form of the data. All bonds will be 50% reduced under these conditions (Fig. 2E), which clearly does not represent the experimental data (Fig. 2A). If the condition is applied that disulfide 5 only forms if disulfide 2 has formed, the total number of disulfide states reduces from 32 to 24 (Fig. 2D) and the percent of reduced disulfides better represents that observed experimentally (Fig. 2E). These simulations indicate that disulfide formation in the β-nodule, and likely elsewhere in the protein, is conditional, which reduces the total number of possible disulfide-bonded states. Nevertheless, there still may be hundreds, if not thousands, of possible disulfide-bonded states of fibrinogen in the circulation.

### A subset of fibrinogen covalent states forms fibrin oligomers

We examined the functional relevance of this biology by determining the disulfide-bonded states of fibrinogen that produces polymerised fibrin. Fibrin forms the scaffold of clots. Formation of fibrin is triggered by thrombin cleavage of fibrinopeptides A and B from the N-termini of the Aα and Bβ chains of fibrinogen (Fig. 3A). Removal of the fibrinopeptides exposes ‘knobs’ that bind to ‘holes’ in the γ- and β-nodules to initiate fibrin fibre formation (Fig. 3A).

**Fig. 3.**
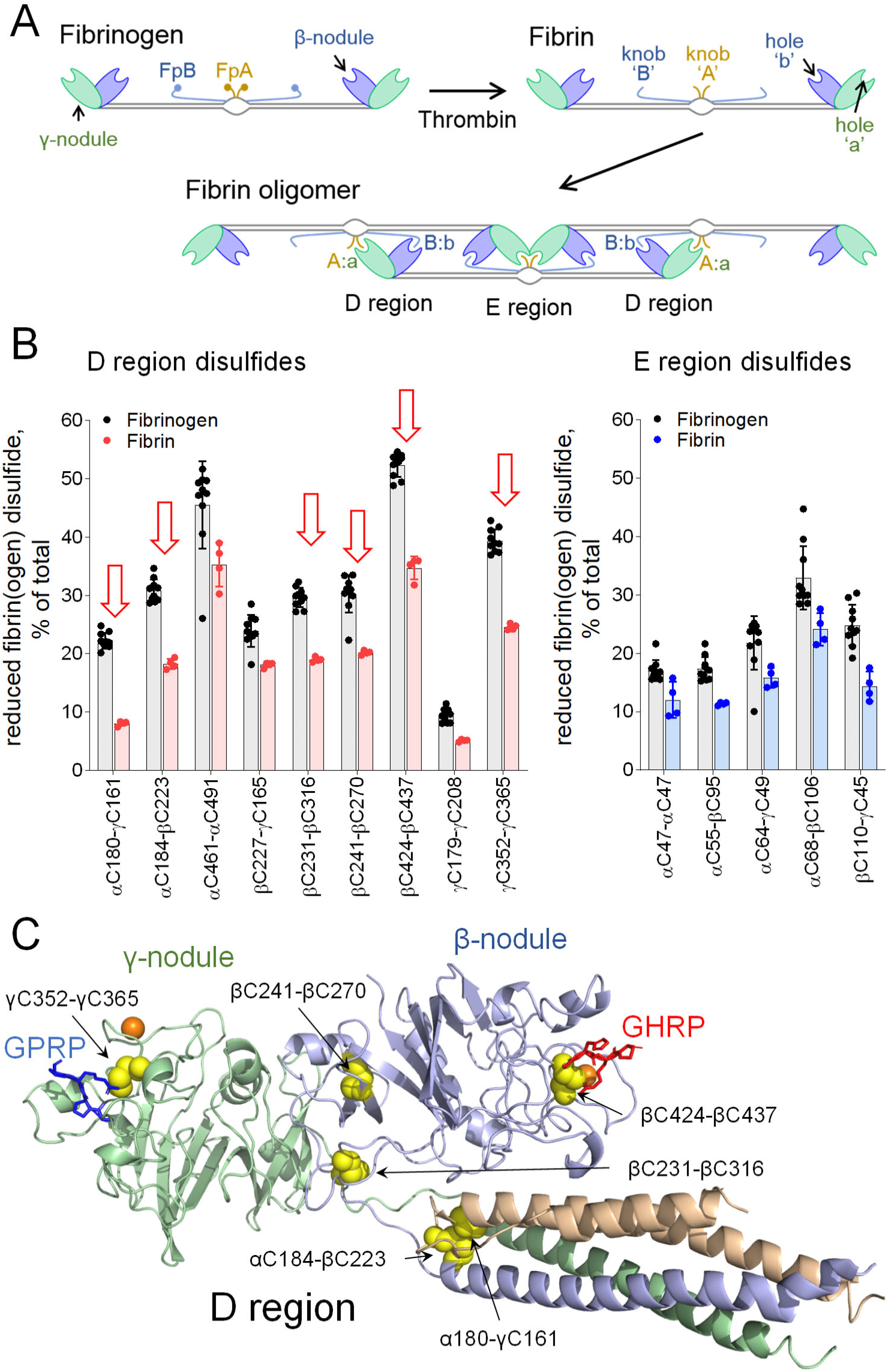
A subset of fibrinogen covalent states forms fibrin oligomers. **A**. Conversion of fibrinogen to fibrin is mediated by thrombin proteolysis of fibrinopeptides (Fp) A and B in the central E region to form knobs ‘A’ and ‘B’. Fibrin oligomers are formed when the exposed knobs ‘A’ and ‘B’ bind to holes ‘a’ and ‘b’ in the γ- and β-nodules, respectively. **B**. Redox states of the nine D region and five E region disulfides in fibrinogen (10 donors) versus fibrin (4 donors). The bars and errors are mean ± SD. Major differences in the redox state of disulfides in fibrinogen versus fibrin are indicated by red arrows. **C**. Ribbon structure of the fibrin D region^33^ with peptide mimetics (sticks) bound in the ‘a’ (GPRP) and ‘b’ (GHRP) holes. The α chains are in wheat, β chains in light blue and γ chains in light green. The positions of the 6 disulfide bonds (yellow spheres) that are significantly more oxidized in fibrin than in fibrinogen are indicated by yellow spheres. Bound calcium ions are shown as orange balls.

Fibrin was produced in healthy human donor plasma by adding thrombin. The fibrin polymer was collected on a plastic rod and the redox state of the fibrin disulfides quantified and compared to the redox state of the plasma fibrinogen pool. Six of the nine D region disulfides in fibrin were >30% more oxidized than the same bonds in plasma fibrinogen, while there was little difference in the redox state of the five E region disulfides in fibrin and fibrinogen (Fig. 3B). This result indicates that a subset of fibrinogen disulfide-bonded states containing more oxidized γ- and β-nodules is preferentially used for fibrin fibre formation.

This may relate to preference for more rigid disulfide-bonded D region nodule holes. For instance, the γC352-γC365 and βC424-βC437 disulfide bonds line the γ- and β-nodule knob-binding pockets, respectively, and adjacent calcium-binding pockets (Fig. 3C). Calcium binding promotes lateral aggregation of fibrin polymers^14^ and reinforces the γ-nodule hole against mechanical forces^15^. Rigid disulfide-bonded binding pockets may be required to withstand the mechanical stresses associated with formation of a fibrin mesh^16^. We cannot rule out, however, that formation of fibrin polymers triggers disulfide formation in the β-nodules.

### The covalent states of fibrinogen are changed by external stimuli

An important question is whether the different fibrinogen states are static or dynamic, that is can they change in their native environment. Cleavage of disulfide bonds is a highly directional chemistry that is influenced by distortion of the protein in which the bond resides^17, 18, 19^. Mechanical strain changes the alignment of the sulfur atoms involved in the cleavage and alters the rate of bond cleavage^20, 21^. Fibrinogen is subjected to mechanical shear forces in the circulation, so we tested the effect of this external stimuli on the covalent states of the protein.

Healthy donor plasma was subjected to fluid shear forces found in arterioles (2000 s^-1^) and stenotic blood vessels (10000 s^-1^)^22^. The 2000 s^-1^ shear rate did not change the redox state of the 14 fibrinogen disulfides, while the pathological fluid shear caused >20% reduction of all 14 disulfides (Fig. 4A). The high shear, therefore, triggered global cleavage of fibrinogen disulfides, indicating that the bonds are dynamic in plasma.

**Fig. 4.**
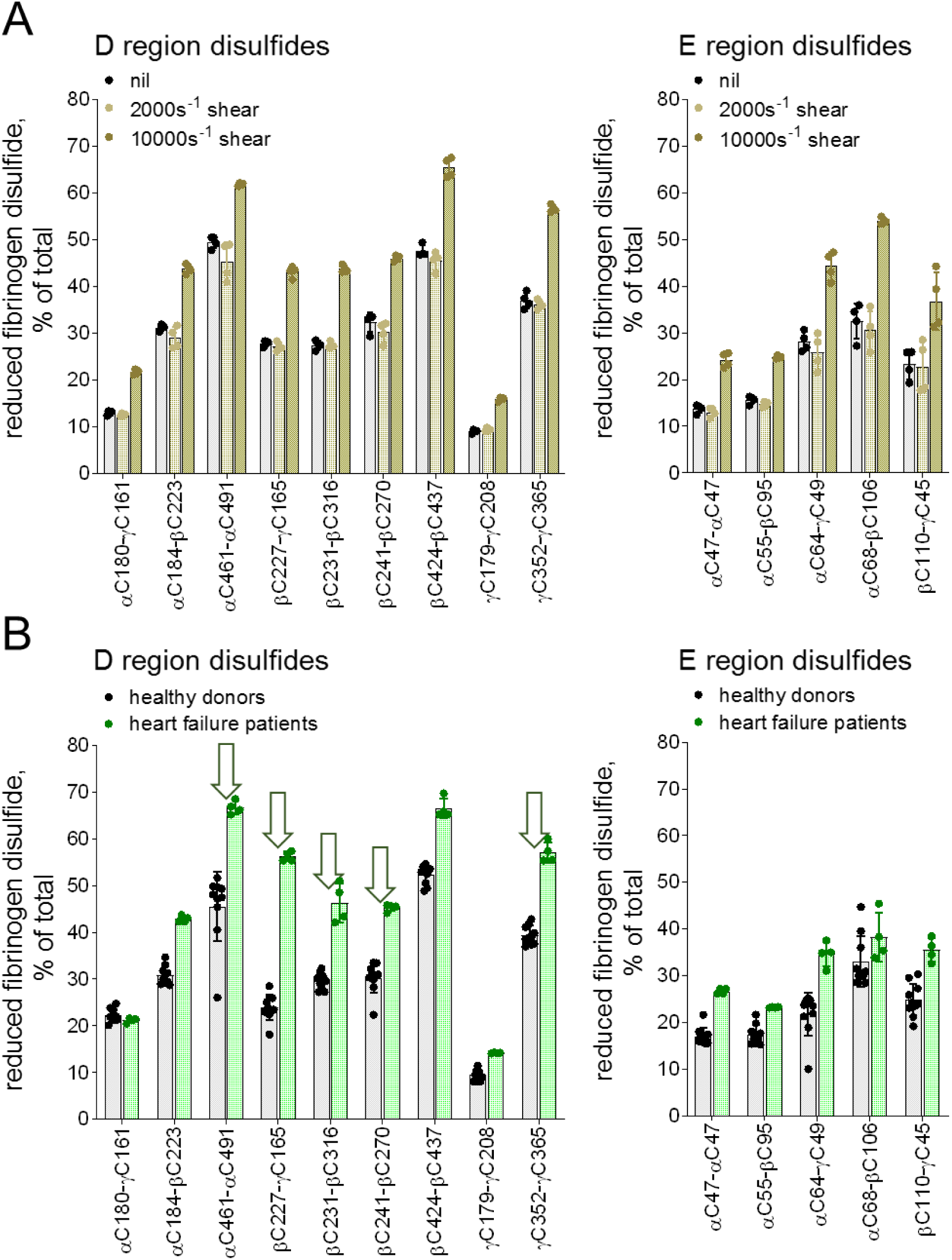
The covalent states of fibrinogen are changed by external stimuli and altered in patients with heart failure. **A**. Redox states of the nine D region and five E region fibrinogen disulfides of four different healthy donor plasmas sheared at rates of 2000s^-1^ or 10000s^-1^ for 5 min. The bars and errors are mean ± SD. **B**. Redox states of the nine D region and five E region fibrinogen disulfides in healthy donors (n = 10) versus patients with severe congestive cardiac failure (n = 4). The bars and errors are mean ± SD. Major differences in the redox state of fibrinogen disulfides are indicated by green arrows.

### The covalent states of fibrinogen are altered in patients with heart failure

The dynamic nature of the fibrinogen disulfides suggested that they may be altered in patients with circulatory disease. Heart failure is a circulatory disease and common independent risk factor for venous thromboembolism^23^. Thromoboprophylaxis is usual for high-risk patients. We examined whether the fibrinogen states were altered in patients with severe congestive cardiac failure.

Five of the nine D region disulfides were >30% more reduced in plasmas of patients than the same bonds in healthy donor plasma fibrinogen (Fig. 4B), while there was little difference in the redox state of the five E region disulfides. This result indicates that heart failure precipitates a selective reduction of some of the D region disulfides in circulating fibrinogen, in contrast to the general reduction when the fibrinogen is subjected to high mechanical shear. The selective dynamic nature of the covalent states of fibrinogen further support the functional relevance of this biology.

### von Willebrand factor also exists in multiple covalent states in the circulation

To test whether the multiple disulfide-bonded states of fibrinogen is a property of other circulating proteins, we examined von Willebrand factor (VWF). While circulating fibrinogen is produced largely by the liver, although it is expressed in many organs, VWF is produced by vascular endothelial cells and megakaryocytes. VWF functions in hemostasis and thrombosis by tethering platelets to the injured blood vessel wall and chaperoning blood coagulation cofactor factor VIII^24^.

VWF circulates as a series of multimers containing variable numbers of dimeric units. The monomer consists of several modular domains and contains a predicted 29 disulfide bonds. The redox state of 13 disulfide bonds in the TIL3, D3 and D4 domains was quantified. The TIL3-D3 disulfides have been structurally defined^25^, while the D4 disulfides are predicted from homology to the D3 domain (Fig. 5A). As we observed for fibrinogen, VWF also exists as multiple disulfide-bonded states in the circulation. The 13 disulfides in the TIL3, D3 and D4 domains ranged from 3 to 30% reduced in 10 healthy human donors (Fig. 5B).

**Fig. 5.**
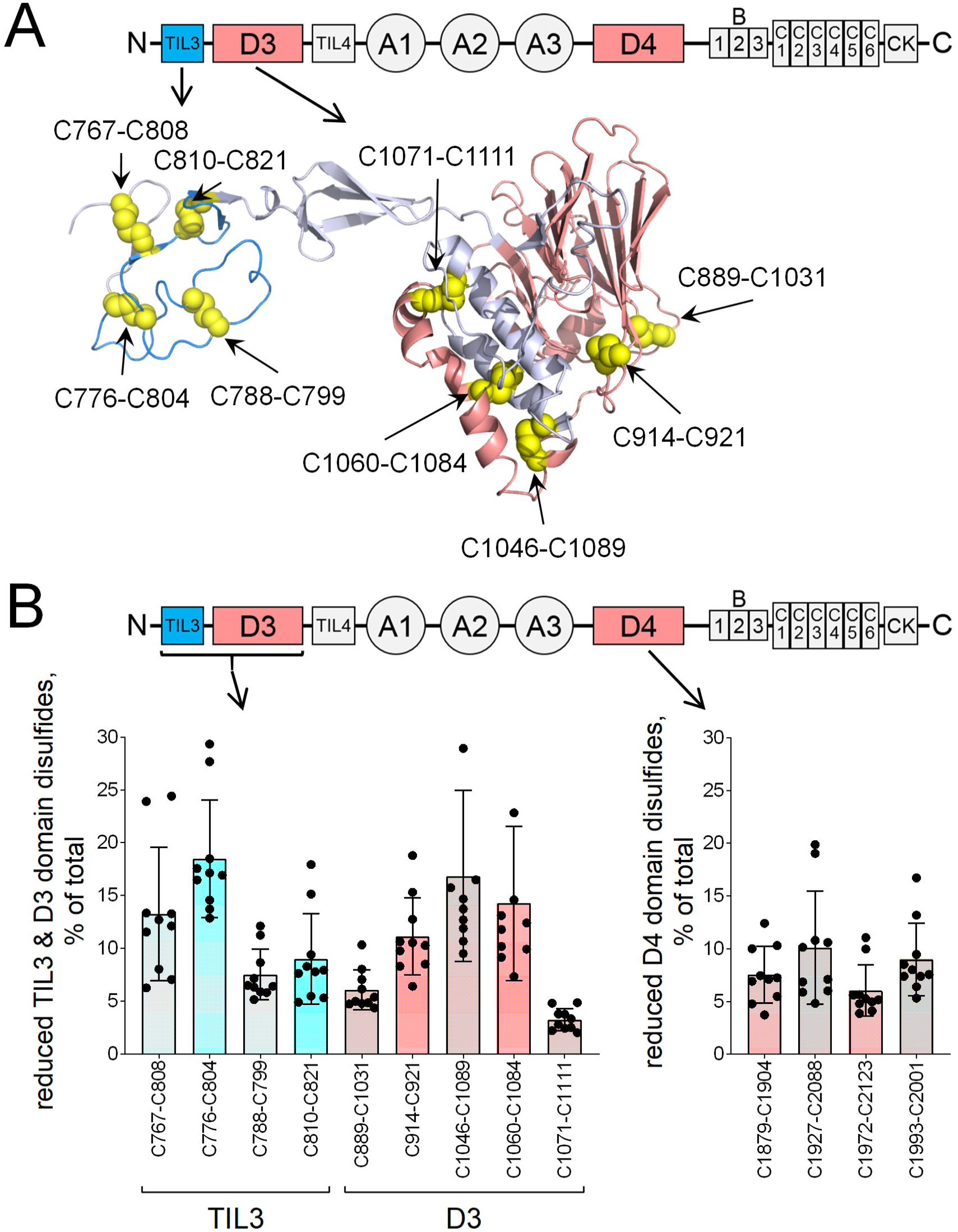
von Willebrand factor also exists in multiple covalent states in the circulation. **A**. Domain structure of VWF is shown at top. Below is the ribbon structure of the TIL3-D3 region^25^ showing the nine disulfide bonds (yellow spheres). The disulfide pairing of the D4 domain is predicted from homology to the D3 domain. **B**. Redox states of the nine VWF TIL3-D3 domain disulfides and four D4 domain disulfides in ten healthy human donors. The bars and errors are mean ± SD.

## Discussion

Our findings demonstrate that the polypeptide backbones of fibrinogen and VWF exist not in a single covalent form in the circulation but rather many different covalent forms. The different covalent forms are interchangeable under certain conditions and, for fibrinogen, some of the forms are preferentially used for fibrin formation. These results have potentially general implications for our understanding of circulating protein function. It is highly likely that fibrinogen and VWF are the first examples of many proteins that exist in different disulfide-bonded states. Our preliminary findings indicate that the protease inhibitor, α2-macroglobulin, and the zymogen, prothrombin, also exist in multiple covalent states in the circulation. These plasma proteins were studied initially because of their relative abundance and ease of capturing the different states in their native environment. It will be interesting to determine whether membrane proteins of circulating leukocytes and platelets, for instance, behave in the same way.

A compelling question is why have these plasma proteins evolved to exist in multiple disulfide-bonded states? One possibility is to facilitate their folding and maturation. Protein folding is a spontaneous process that is dominated by the second law, ΔG = ΔH - TΔS, where a net negative Gibbs free energy (G) is achieved through weighing of the enthalpy (ΔH) and entropy (-TΔS) contributions. The conformational entropies of the more reduced disulfide-bonded states of fibrinogen and VWF are predicted to be higher than the more oxidized states. This may mitigate to some extent the negative entropic effects of folding. It is possible that these entropic gains were a driving force for the evolution of these protein states.

A striking feature of the different covalent states of fibrinogen is how little the balance of the states differs from donor to donor. It is not appreciably influenced by donor age or gender. The results for VWF demonstrate more donor-to-donor variation than fibrinogen, although there is a consistent pattern in the redox states of the disulfide bonds. These findings suggest that the mix of the forms is tightly controlled, either during maturation of the protein in the cell and/or following secretion into the circulation. One class of enzymes that may be involved are the oxidoreductases that manipulate protein disulfide bonds in the ER/Golgi^26^ and in the circulation^27^.

The implications of our findings for thrombophilia’s, drug resistance and drug development are manifold. Perturbation of the mix of the covalent forms of fibrinogen and VWF, and other circulating proteins, could tip the balance between bleeding and clotting in individuals and underlie some hereditary or acquired thrombophilia’s^28^. In addition, anti-thrombotic therapeutics are commonly associated with drug resistance^29^. If a drug binds preferentially to one or more covalent forms of a protein, then events that reduce the incidence of these forms in individuals would result in drug resistance. Similarly, if drugs are developed against some covalent forms of a protein target but not others, this will inevitably result in drug resistance in some individuals. In experimental settings, the multiple covalent states of a protein should be considered when these proteins are investigated. Antibodies and small molecule inhibitors may efficiently bind and inhibit some of the covalent states but not others.

It was recognized 75 years ago by Erin Schrӧdinger that proteins have enough potential variety in their configurations to encode huge amounts of information^30^. Existing in multiple disulfide-bonded states is an effective and efficient way of maximising the variety of information a single protein can convey.

## Methods

All procedures involving collection of human blood from healthy volunteers were in accordance with the Human Research Ethics Committee of the University of Sydney (HREC 2014/244), Alfred Hospital Ethics, Monash University Standing Committee for Research in Humans, and the Helsinki Declaration of 1983. Blood from 14 healthy donors on no medications (7 male, 7 female, 22-58 years old) and four patients with cardiac comorbidities including severe congestive cardiac failure (1 male, 3 female, 29-59 years old) was drawn into ACD-A tubes (BD Vacutainer) and plasma collected by twice centrifugation at 800 g for 20 min at room temperature. On some occasions, plasma was sheared at rates of 2000 s^-1^ or 10000 s^-1^ for 5 min at room temperature using a Kinexus pro+ rheometer. Plasma (0.7 mL) was incubated with polyclonal anti-fibrinogen or anti-VWF antibody-coated (Dako) Dynabeads (Life Technologies) on a rotating wheel for 1 h at 22°C. The beads were collected, excess plasma aspirated, and incubated in 0.3 mL of 5 mM ^12^C-IPA in phosphate-buffered saline containing 10% DMSO for 1 h at 22°C in the dark to alkylate unpaired Cys thiols in the proteins. The supernatants were aspirated, and the beads incubated with NuPAGE LDS sample buffer (Life Technologies) containing a further 5 mM ^12^C-IPA for 30 min at 60°C. The second labelling step was to ensure complete alkylation of Cys residues. The supernatants were resolved on SDS-PAGE.

Fibrin was prepared in 1 mL of plasma by addition of 50 units of bovine thrombin (Sigma, SRE0003) and 10 mM CaCl_2_ and MgCl_2_ for 22°C. The fibrin clots were collected using a plastic rod, blotted with tissue paper, and submerged in 1 mL of 5 mM ^12^C-IPA in phosphate-buffered saline containing 10% DMSO for 1 h at 22°C in the dark. The fibrin was washed 3 times with phosphate-buffered saline before dissolving the polymer in 1 mL of 20 mM acetic acid for 24 h at 22°C. Twenty microliters of the solution was incubated with NuPAGE LDS sample buffer (Life Technologies) containing 5 mM ^12^C-IPA for 30 min at 60°C and the proteins resolved on SDS-PAGE.

The SDS-PAGE gels were stained with colloidal coomassie (Sigma) and the fibrin(ogen) or VWF bands excised, destained, dried, incubated with 40 mM dithiothreitol and washed^12^. The gel slices were incubated with 5 mM ^13^C-IPA (Cambridge Isotopes) for 1 h at 22°C in the dark to alkylate the disulfide bond Cys, washed and dried before digestion of proteins with 12.5 ng/μl of trypsin (Roche) in 25 mM NH_4_CO_2_ and 10 mM CaCl_2_ overnight at 25°C. Peptides were eluted from the slices with 5% formic acid, 50% acetonitrile. Liquid chromatography, mass spectrometry and data analysis were performed as described^31, 32^. Cys labelled with ^12^C-IPA or ^13^C-IPA has a mass of 133.05276 or 139.07289, respectively. Thirty Cys-containing peptides were analysed (**Table 1**). The different redox forms of the Cys residues were quantified from the relative ion abundance of peptides labelled with ^12^C-IPA and/or ^13^C-IPA.

## Acknowledgments

We thank Elizabeth Gardiner, Amanda Davis, Deirdre Murphy and Robert Andrews for the heart failure patient plasmas, and Aster Pijning and Manuela Florido for helpful discussions.

## Author contributions

PJH conceived the study, designed experiments and wrote the manuscript. DB designed and performed the experiments

## Competing interests

Authors declare no competing interests.

## Supplementary Information

**Table S1.**
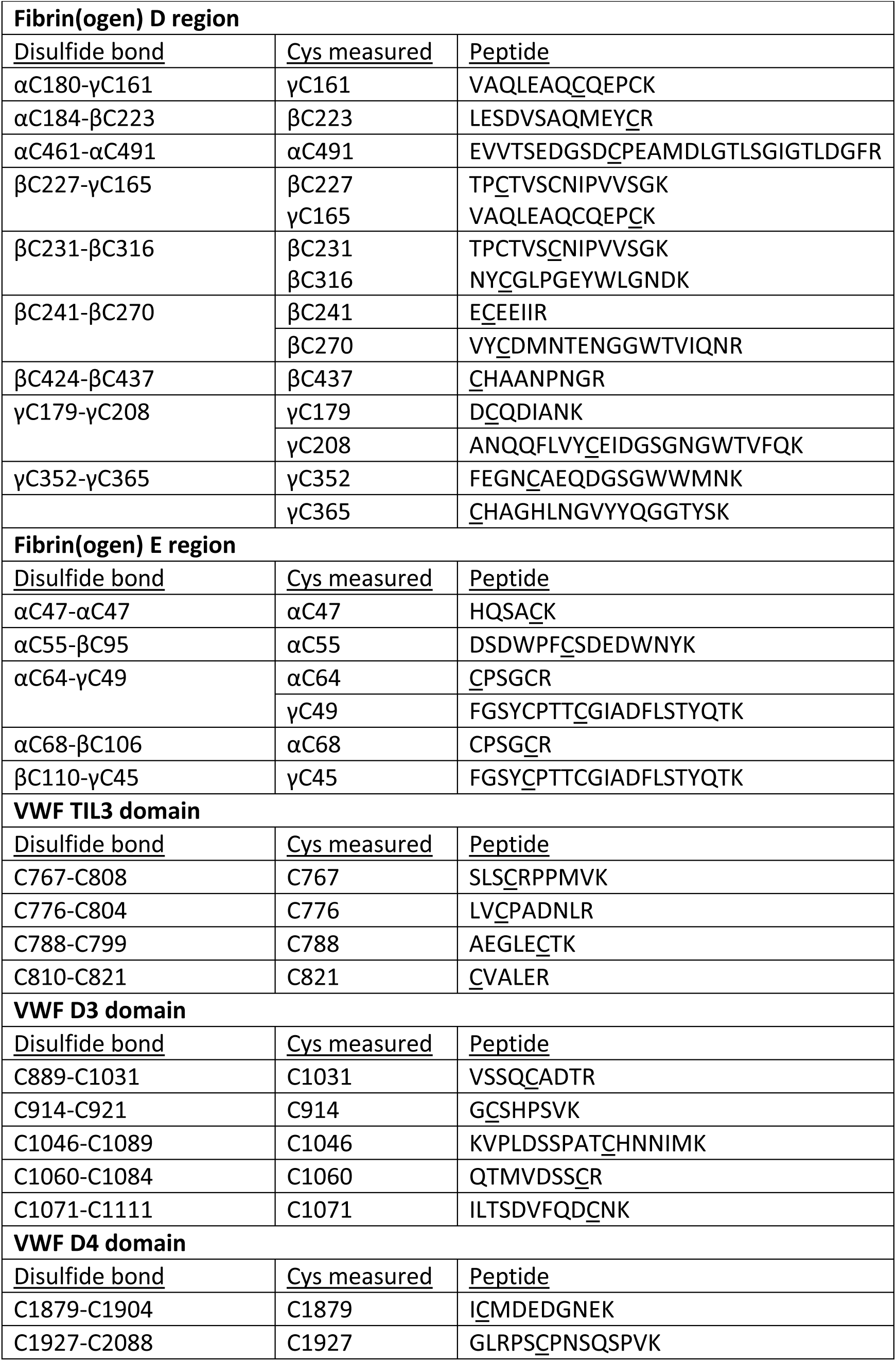
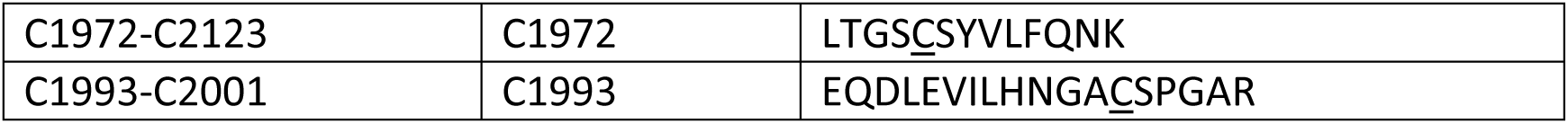
Cysteine containing peptides analysed by HPLC and mass spectrometry. Cysteine numbering is according to UniProt identifiers P02671, P02675 and P02679 for human fibrinogen α, β and γ chains, respectively, and P04275 for human von Willebrand factor. The Cys residues of the disulfide that were measured are underlined in the peptide.

**Figure S1.**
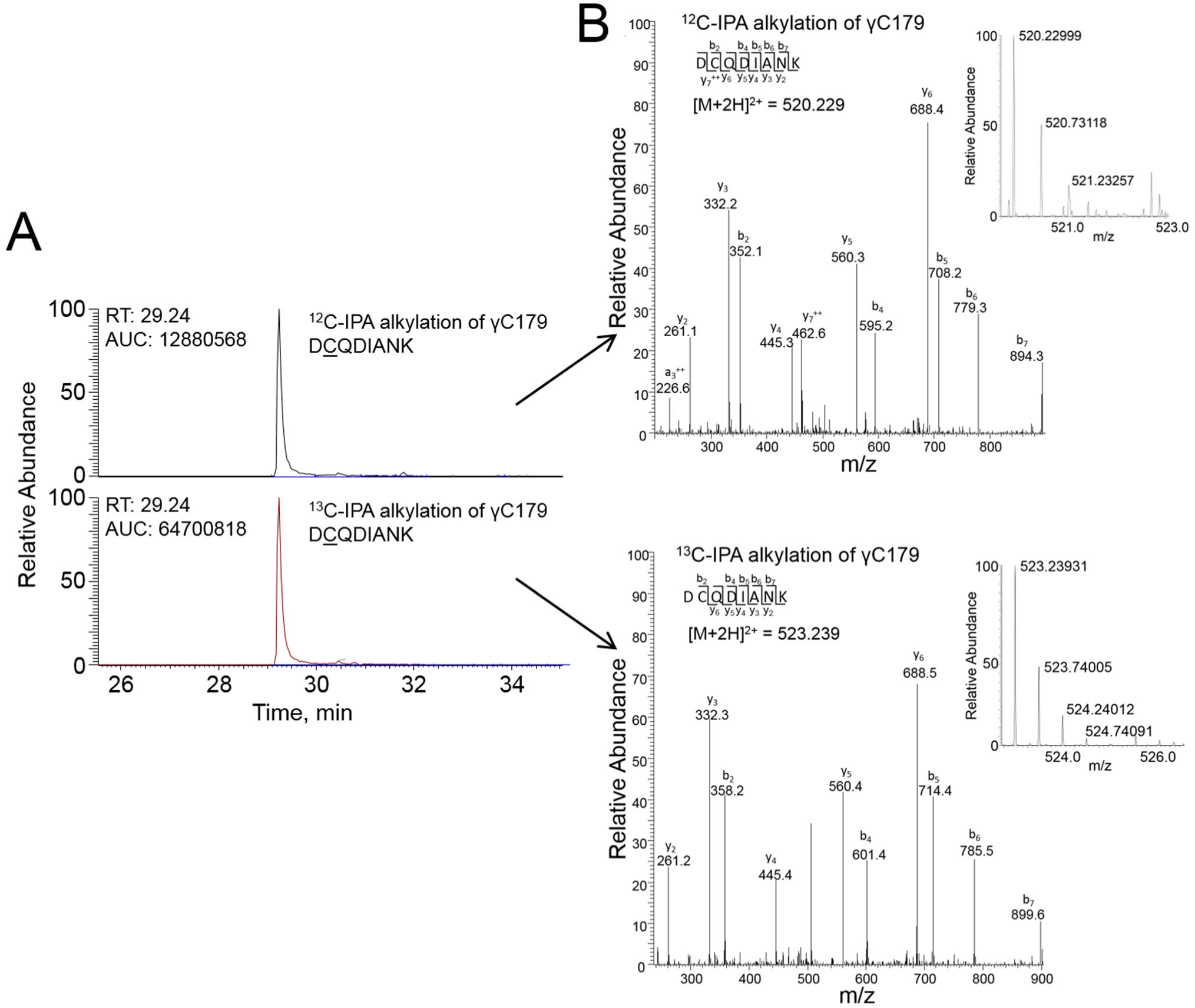
Differential cysteine alkylation of the fibrinogen γ chain Cys179 residue and peptide analysis. **A**. HPLC resolution of the γ chain D<u>C</u>QDIANK peptide containing Cys179 labelled with either ^12^C-IPA (upper trace) or ^13^C-IPA (lower trace). **B**. Representative tandem mass spectra of the γ chain D<u>C</u>QDIANK peptide. The upper and lower traces are examples of ^12^C-IPA or ^13^C-IPA alkylation of Cys179, respectively. The accurate mass spectrum of the peptide is shown in the insets (upper trace, observed [M+2H]^2+^ = 520.229 m/z and expected [M+2H]^2+^ = 520.229 m/z; lower trace, observed [M+2H]^3+^ = 523.239 m/z and expected [M+2H]^2+^ = 523.239).

